# Unleashing the Immune Arsenal: Development of Broad-spectrum Multiepitope Bluetongue Vaccine Targeting Conserved T Cell Epitopes of Structural Proteins

**DOI:** 10.1101/2024.04.12.589199

**Authors:** Harish Babu Kolla, Anuj Kumar, Mansi Dutt, Roopa Hebbandi Nanjunadappa, Karam Pal Singh, Peter Paul Clement Mertens, David Kelvin, Channakeshava Sokke Umeshappa

## Abstract

Bluetongue (BT) is a severe arboviral disease affecting sheep, cows, and other wild ruminants, caused by the Bluetongue virus (BTV). The virus has evolved into over 32 serotypes, rendering existing vaccines less effective. While the structural proteins of this virus represent promising targets for vaccine development, they unfortunately exhibit high amino acid polymorphism and are laden with numerous inhibitory epitopes. Structural proteins such as VP1 and VP7 are highly conserved and may contain epitopes capable of triggering cross-reactive cell-mediated immunity (CMI). In this study, we identified highly conserved MHC-I and -II-restricted T cell epitopes within VP1, VP5, and VP7 BTV proteins and developed an effective *in silico*-immuno-informatics-based broad-spectrum BT multiepitope vaccine for bovine and laboratory mouse systems. The conserved epitopes utilized in the vaccines are highly antigenic, non-allergenic, non-toxic, and capable of inducing IFN-*γ* (only CD4+ T cell epitopes). Both mouse and bovine vaccines were tethered with Toll-like receptor (TLR)-4-agonist adjuvants, beta-defensin 2, and the 50s ribosomal unit to stimulate innate immunity for CMI development. Protein-protein docking analysis revealed strong binding affinities, while extensive 100-nanosecond molecular dynamics simulations indicated stable complexes between the vaccine structures and TLR4. Vaccination simulation studies demonstrated their ability to trigger proinflammatory responses. Therefore, these novel vaccine designs necessitate further exploration through wet lab experiments to evaluate their immunogenicity, safety, and effectiveness for practical deployment in livestock.

## Introduction

Bluetongue (BT) disease is widely distributed around the world and is one of the most economically important arboviral diseases of ruminants (primarily sheep). It is caused by bluetongue virus (BTV), belonging to genus *Orbivirus* within a new family *Sedoreoviridae* and the order *Reovirales* (Mertens 2004, Matthijnssens 2022). BTV genome comprises 10 double-stranded RNA segments of various lengths (Seg-1 to Seg-10 in decreasing order of size), which encodes for the various proteins, such as Viral protein (VP) 1, VP2, VP3, VP4, VP5, VP6/6A, VP7 and Non-structural (NS) protein 1, NS2, and NS3/3A (Verwoerd and Huismans 1972, Maan 2012). Conventional Bluetongue virus (BTV) serotypes are transmitted via the bites of blood-feeding midges belonging to the *Culicoides* genus (*Ceratopogonidae* family). This infection induces severe damage to vascular endothelial cells, leading to edema, hemorrhage, vascular thrombosis, and tissue infarction (MacLachlan 1994, Umeshappa, Singh et al. 2011). Animals affected by BTV typically experience lameness, weakness, reduced productivity, and varying degrees of mortality, resulting in considerable economic losses (MacLachlan 1994). BTV can persist in the bloodstream for an extended period, raising concerns about the movement of animals across borders (MacLachlan 1994).

Currently, there are more than 32 reported serotypes of Bluetongue virus (BTV) globally, including eight recently identified ‘novel’ types that may not infect vector insects. These newly emerged serotypes have the potential to exhibit unpredictable behavior, differing from the established 24 serotypes. The presence and widespread distribution of multiple BTV serotypes, along with the constant emergence, reemergence, and co-circulation of various serotypes, pose challenges to the effectiveness of existing vaccines, such as live attenuated vaccines (LAV) and inactivated vaccines (IAVs), which are predominantly serotype-specific (Rojas 2021). DNA plasmid-vectored vaccines have shown efficacy in mouse models (Calvo-Pinilla E 2009, Jabbar TK 2013), their practical deployment faces hurdles, including the risk of affecting genes responsible for cell growth, reduced immunogenicity, and high production costs for large-scale manufacturing. Recent studies on broad-spectrum influenza vaccines have highlighted the potential of targeting conserved immunodominant epitopes within viral proteins to develop pan-BTV vaccines effectively (Xie 2019, De Jong, Aartse et al. 2020, Lo, Misplon et al. 2021). Consequently, we have developed an innovative *in silico* immunoinformatics-based pan-BTV vaccine, utilizing conserved epitopes from non-structural proteins, NS1 and NS2 (Harish Babu Kolla 2023). This vaccine design has potential to induce cross-protective cell-mediated immune (CMI) responses against all BTV serotypes. In this study, we aim to investigate the feasibility of developing a similar pan-BTV vaccine by targeting the structural proteins of BTV.

Similar to NS proteins, certain structural proteins, notably VP1 and VP7, exhibit higher amino acid sequence conservation (Maan, Maan et al. 2010, Belaganahalli 2011, Belaganahalli 2012), suggesting their potential to harbor conserved T cell epitopes. Thus, in this investigation, we employed in silico methods, such as multiple sequence alignments and epitope mapping, to analyze the key structural proteins—VP1, VP2, VP5, and VP7—of all the BTV serotypes, sourced from the National Center for Biotechnology Information (NCBI) database. We specifically analyzed these proteins for the presence of conserved T cell epitopes recognized by murine Major Histocompatibility Complex (MHC) and Bovine Leukocyte Antigen (BoLA) molecules. VP3 was excluded from our analysis due to documented inhibitory effects on immune response development (Fablet, Kundlacz et al. 2022). Given the substantial variability among serotypes, the outer capsid of the virus—comprised of VP2 and VP5— is shown to induce serotype-specific neutralizing antibodies (Jeggo, Wardley et al. 1984, Rojas, Rodriguez-Calvo et al. 2017). As a result, B cell epitopes have been deliberately omitted from our broad-spectrum vaccine design. Instead, we focused on eliciting cell-mediated immune (CMI) responses, as they have demonstrated cross-protection against multiple BTV serotypes in both infections and vaccinations of sheep (Jeggo, Wardley et al. 1984, Takamatsu 1989, Calvo-Pinilla E 2009, Umeshappa, Singh et al. 2011, Rojas, Rodriguez-Calvo et al. 2017). Notably, viral vectors expressing other BTV proteins such as NS2, VP2, and VP7 have shown to confer CD8+ T cell-mediated cross-protection against various BTV serotypes (Jones 1997, Bouet-Cararo, Contreras et al. 2014, Martín 2015). Moreover, studies by Roja et al. elegantly demonstrated the relevance of incorporating epitopes from BTV proteins in vaccine formulations (Rojas, Rodriguez-Calvo et al. 2011, Rojas 2014). We selected the mouse model for its increasing utility in studying Bluetongue (BT), including assessing vaccine efficacy, while bovines were chosen due to their status as natural hosts susceptible to BTV infection, akin to sheep.

In this study, we identified the conserved CD8+ and CD4+ T cell epitopes with high antigenicity, non-allergenicity, non-toxicity, and the ability to induce IFN-gamma (IFNγ) in the VP1 and VP7 proteins corresponding to the murine MHC alleles and bovine BoLA alleles. These epitopes along with TLR4 agonists were utilized in the development of *in silico*-based pan-BTV vaccines. Protein-protein docking analysis demonstrated robust binding affinities, and extensive 100-nanosecond molecular dynamics simulations confirmed the stability of the complexes formed between the vaccine structures and TLR4. Immune response simulation studies revealed vaccines’ capacity to trigger an immune response. This comprehensive study serves as a proof of concept for future research efforts focused on creating potent pan-BTV vaccines. These vaccines have the potential to offer cross-protection against all known BTV serotypes, thereby mitigating the dissemination of these virulent strains and control of BT.

## Methodology

### Sequence retrieval and multiple sequence alignment

Firstly, we mined the full length amino acid sequences of the BTV structural proteins, such as VP1, VP2, VP5 and VP7, from all the 24 serotypes that are available in NCBI taxonomy browser (https://www.ncbi.nlm.nih.gov/Taxonomy/Browser/wwwtax.cgi?id=40051). Later, we aligned the amino acid sequences of all the 4 proteins using BLOSUM 65 matrix of ClustalW module in the Molecular evolutionary genetic analysis (MEGA-X) software. The alignment files of all the proteins were saved in MEGA format for further analysis. Then the amino acid sequence of all the proteins was determined using the MEGA files.

### Prediction of murine MHC I- and II-restricted epitopes

We predicted the MHC class I and II specific CD8+ and CD4+ T cell epitopes in all the 4 proteins that are considered for the analysis. Later, the conserved T cells were identified based on their amino acid sequence conservation among all 24 BTV serotypes. For epitope prediction, we used the amino acid sequences of VP1, VP2, VP5 and VP7 proteins from BTV-1 serotype. Prediction of MHC class I alleles (H2-Kb and H2-Db) in C57BL/6 mice was performed utilizing the “NetMHCpan BA 4.1” module available on the Immune Epitope Database Epitope Analysis Resource (IEDB-AR) website (http://tools.iedb.org/mhci/ (Vita, Mahajan et al. 2019). The host was selected as mouse for the epitope prediction. Epitope selection relied on Half-maximal Inhibitory Concentration (IC50) values, with a threshold of <500nM set for CD8+ T cell epitopes (Nielsen, Lundegaard et al. 2003). Likewise, predictions for CD4+ T cell epitopes were conducted for the MHC II H2-IA^b^ allele in C57BL/6 mice, employing the “MHCIIpan 4.0 BA” module accessible on the IEDB-AR tool website (http://tools.iedb.org/mhcii/)(Vita, Mahajan et al. 2019). Threshold parameters for CD4+ T cell epitopes were set using IC50 values, with <1000nM being considered, which indicates the strong interaction between the epitope peptide and the MHC allele (Southwood 1998, Sidney, Steen et al. 2010). The predictions focused on the VP1, VP2, VP5 and VP7 structural proteins of the BTV1 serotype genome.

### Bovine BoLA class I- and II-restricted epitope prediction

Likewise, we predicted the CD8+ and CD4+ T cell epitopes in the VP1, VP2, VP5 and VP7 proteins corresponding to the bovine immune system. The prediction was carried out utilizing the IEDB-AR server (http://tools.iedb.org/mhci/) (Dhanda, Mahajan et al. 2019), with the host species specified as “cow.” While predicting the CD8+ T cell epitopes, the most frequent BoLA class I alleles such as BoLA-BoLA-1*02301, BoLA-2*01201, BoLA-3*00201, BoLA-4*02401, BoLA-6*01301 and BoLA-6*01302 were selected (Pandya, Rasmussen et al. 2015, Svitek 2015). For predicting CD4+ T cell epitopes, we used NetBOLAIIPan 1.0 (https://services.healthtech.dtu.dk/service.php? NetBoLAIIpan-1.0) (Fisch, Reynisson et al. 2021). These alleles have previously been employed for predicting BoLA-restricted class I epitopes for foot-and-mouth virus, which shares similarities with BT during the early stages of infection (Williamson S 2008). A threshold IC50 value of <500nM was utilized for epitope prediction. The CD4+ T cell epitopes were predicted corresponding all the BoLA II alleles that were deposited in the tool (BoLA-BoLA-DRB3_0101, BoLA-DRB3_1001, BoLADRB3_1101, BoLA-DRB3_1201, BoLA-DRB3_1501, BoLADRB3_1601 and BoLADRB3_2002). The tool generates potential predicted T cell epitopes in 15 amino acid length, each with different %Rank EL scores. A threshold parameter of %Rank EL scores less than 1.0 was utilized for the prediction of CD4+ T cell epitopes (Fisch, Reynisson et al. 2021).

### Analysis of the epitopes for pan-BTV vaccine development

To obtain conserved CD8+ and CD4+ T cell epitopes, we examined the predicted epitopes within the VP1, VP2, VP5 and VP7 proteins of BTV1 by comparing them with amino acid alignment files previously generated using MEGA-X software. Our analysis aimed to determine the extent of conservation of these epitopes within the alignment files encompassing VP1, VP2, VP5 and VP7 proteins across all 24 serotypes. The conserved T cell epitopes were further screened before designing a multi-epitope vaccine. These epitopes were subjected for screening based on their antigenicity, allergenticity, toxicity and their ability to induce IFNg. Firstly, the antigenicity of these epitopes was predicted using VaxiJen v2.0 server (http://www.ddg-pharmfac.net/vaxijen/VaxiJen/VaxiJen.html) (Doytchinova and Flower 2007). Similarly, allergenicity and toxicity of these epitopes was predicted using online resources called AllerTOP v2.0 (https://www.ddg-pharmfac.net/AllerTOP/) (Dimitrov, Flower et al. 2013, Dimitrov, Bangov et al. 2014) and ToxinPred (https://webs.iiitd.edu.in/raghava/toxinpred/algo.php) (Gupta, Kapoor et al. 2013). Finally, the INFg inducing potentail of the epitopes was determined with the help of IFNepitope tool (https://webs.iiitd.edu.in/raghava/ifnepitope/application.php) (Dhanda, Vir et al. 2013).

### Designing pan-BTV vaccine

Following a rigorous epitope screening process, those epitopes that exhibited antigenicity, non-allergenicity, non-toxicity, and IFN-*γ*-inducing properties were selected for the development of a pan-BTV multi-epitope vaccine capable of eliciting an immune response across all serotypes. During vaccine design, the validated linkers (El Bissati, Chentoufi et al. 2016) were utilized to connect the CD8+ T cell epitopes using the ‘AAY’ linker and CD4+ T cell epitopes with the ‘GPGPG’ linker as previously described by us (Khairkhah, Bolhassani et al. 2022). To enhance the effectiveness of our vaccines, we strategically incorporated an adjuvant sequence at the N-terminal end of the vaccine sequence using the ‘EAAAK’ linker (Ryoichi Arai 2001). We employed two TLR4-activating adjuvants, beta-defensin 2 and a 50S ribosome subunit, as carriers in our vaccine constructs. These adjuvants were chosen for their capability to induce dendritic cell maturation and initiate a robust cell-mediated immune response *in vivo* (Biragyn, Ruffini et al. 2002, Lee, Shin et al. 2014, Machado and Ottolini 2015, Behmard, Soleymani et al. 2020, Kalita, Padhi et al. 2020, Singh, Thakur et al. 2020, Singh, Jakhar et al. 2020, Kumar, Rathi et al. 2022, Fu, Zong et al. 2023).

### Antigenicity, allergenicity, solubility, and physicochemical properties of the vaccine constructs

We devised a total of 4 vaccine constructs, comprising mouse and bovine vaccines, each incorporating two distinct TLR4 agonist adjuvants. These vaccines underwent evaluation for antigenicity and allergenicity as previously described. Additionally, the solubility and physicochemical properties, such as molecular weight, theoretical isoelectric point (pI), instability index (II), aliphatic index (AI), and grand average of hydropathicity index (GRAVY), were predicted using the Protein-Sol (http://protein-sol.manchester.ac.uk/) (Hebditch, Carballo-Amador et al. 2017) and ProtParam tool (physico-chemical properties) (https://web.expasy.org/protparam/) (Wilkins, Gasteiger et al. 1999), respectively.

### Structure modeling and evaluation

We employed an *in silico* protein modeling approach to predict the three-dimensional structures of the four vaccines. The 3-D structures of the vaccine constructs were modeled using the Robetta server (https://robetta.bakerlab.org/) (Kim, Chivian et al. 2004). Subsequently, these protein models underwent refinement using the GalaxyWEB server (https://galaxy.seoklab.org/) (Ko, Park et al. 2012) and the quality of the models was assessed based on the phi (ϕ) and psi (Ψ) torsion angles using the PROCHECK module ((Laskowski R A 1993) within the protein structure verification server (PSVS) (https://montelionelab.chem.rpi.edu/PSVS/PSVS/).

### Preparation of TLR4 receptor and molecular docking

Molecular docking studies were conducted to assess the interaction between the vaccines and TLR4, a potent inducer of antiviral activity (Suh, Zhao et al. 2009), as described previously. Briefly, the X-ray crystallography-derived structures of mouse TLR4, identified by the PDB ID 4G8A ID, were obtained from the RCSB-PDB. As the crystal structures of bovine TLR4 are unavailable in the PDB, we retrieved modeled structures from the alpha-fold database for bovine TLR4 (Q9GL65). The receptor molecules were prepared for molecular docking by eliminating heteroatoms and water molecules using PyMOL software. Molecular docking studies were performed using the Cluspro server (https://cluspro.bu.edu/publications.php) (Desta, Porter et al. 2020) with chain ‘A’ assigned to receptors and chain ‘B’ to the vaccine constructs or ligands. The binding affinity between TLR4 and vaccine constructs in each docking complex was assessed using the PRODIGY server (https://wenmr.science.uu.nl/prodigy/) (Vangone and Bonvin 2015), while two-dimensional interactions were visualized using the PDBsum tool (https://www.ebi.ac.uk/thornton-srv/databases/pdbsum/) (Laskowski, Jablonska et al. 2018).

### Molecular dynamics simulations

Molecular dynamics (MD) simulations were carried out for all the four docking complexes to investigate their stability at the atomic level, as described previously. Consistent protocols were applied to each complex, employing state-of-the-art MDS techniques with the AMBER99SB-ILDN protein and AMBER94 nucleic force fields (Lindorff-Larsen, Piana et al. 2010) integrated into the GROMACS 2023 package (Abraham 2015) running on high-performance computing (HPC) infrastructure. The complexes were immersed in solvent using the transferable intermolecular potential 3P (TIP3P) water model and underwent energy minimization with LINear Constraint Solver (LINCS) constraint algorithms to achieve neutrality (Hess 1997). Subsequent equilibration steps involved NVT ensemble at 300K and NPT ensemble with the Parinello-Rahman barostat coupling ensembles (Parrinello 1981). Following equilibration, a 100ns MDS was performed for all four docked complexes, a standard duration in MD simulations for multi-epitope vaccine development (Sirohi, Gupta et al. 2022, Hou, Wu et al. 2023, Jyotisha, Qureshi et al. 2023, Zhu, Tan et al. 2023). Trajectories were analyzed post-simulation, and plots were generated using various GROMACS package modules.

## Results

### Amino acid conservation among the structural proteins in BTV serotypes

We analyzed the amino acid sequence conservation of BTV’s structural proteins, including VP1, VP2, VP5, and VP7. Notably, VP1 and VP7 displayed high conservation levels, with sequence similarities of 95.6% and 73.63%, respectively (**Figure 1A**), indicating the presence of conserved epitopes within these proteins. In contrast, VP5 exhibited moderate variability, with an identity of 39.69%, making it the third most variable protein, while VP2 showed the lowest identity at 10.25% (**Figure 1A**), consistent with previous research (Mertens 1984, Shafiq 2013).

**Figure 1.**
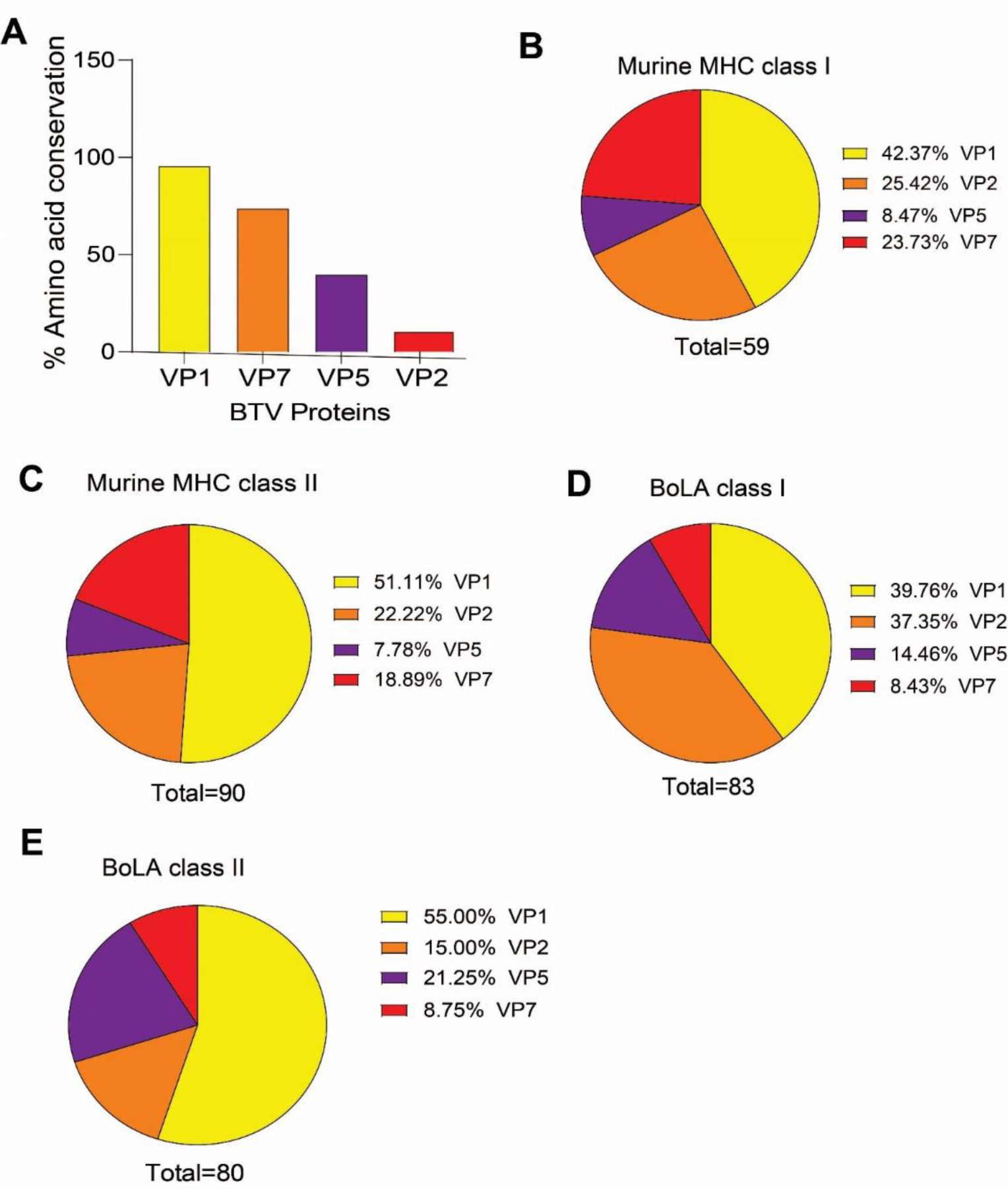
Amino acid sequence conservation in the structural proteins of BTV and distribution of T cell epitopes among them. **A.** Bar graph representing the amino acid sequence conservation among the VP1, VP2, VP5 and VP7 proteins of BTV. The VP1 and VP7 are highly conserved whereas the VP2 and VP5 are more polymorphic. **B-E**. Distribution of murine MHC (B and C) and bovile BoLA (D and E)-restricted CD8+ and CD4+ T cell epitopes in the structural proteins of BTV.

### Frequencies of mouse-specific T cell epitopes in structural proteins of BTV-1

The C57BL/6 mouse model has been widely utilized to investigate the immunopathogenesis of BTV and assess vaccine candidates against the virus. Therefore, we employed this model to establish the proof of concept for our pan-BTV multiepitope vaccines and predicted CD8+ T cell epitopes recognized by H-2-Db and H-2-Kb class I molecules, as well as CD4+ T cell epitopes recognized by H-2-IAb class II molecule. Our predictions yielded a total of 57 CD8+ T cell epitopes for the mouse system, distributed as follows: 25 (42.37 %) in VP1, 15 (25.42 %) in VP2, 5 (8.47 %) in VP5, and 14 (23.73 %) in VP7 proteins of BTV-1 (**Figure 1B, Table S1)**. On the other hand, we identified 90 CD4+ T cell epitopes, with VP1 harboring the majority, accounting for 46 (51.11%) of the epitopes, followed by VP2 with 20 (22.22%), VP5 with 7 (7.78%), and VP7 with 17 (18.89%) epitopes (**Figure 1C, Table S2).** These findings highlight VP1 as the primary source of both mouse CD8+ and CD4+ T cell epitopes.

### Frequencies of bovine-specific T cell epitopes in structural proteins of BTV-1

Like sheep, bovines serve as natural hosts for BTV and are closely related to sheep in terms of genome, immunology, anatomy, and physiology. Unfortunately, tools for predicting T cell epitopes in sheep are not available. However, understanding the nature of BoLA epitope presentation and their conservation can provide insights into T cell epitope presentation in the ovine immune system. Additionally, epidemiological studies emphasize the importance of preventing BT in cattle to curb its spread among cattle and sheep populations. These factors drive our focus on bovines for pan-BTV vaccine development. The IEDB tool contains information on hundreds of BoLA class I alleles. Therefore, we considered the most frequent BoLA class I alleles, including BoLA-6*01301, BoLA-6*01302, BoLA-2*01201, BoLA-4*02401, BoLA-3*00101, and BoLA-1*02301, for CD8+ T cell epitope prediction. We identified a total of 83 CD8+ T cell epitopes, with 33 (39.75%) in VP1, 31 (37.34%) in VP2, 12 (14.45%) in VP5, and 7 (8.43%) in VP7 proteins of BTV1 (**Figure 1D, Table S3)**. For CD4+ T cell epitopes, we found 44 (55%) in VP1, 12 (15%) in VP2, 17 (21.25%) in VP5, and 7 (8.75%) in VP7 of the BTV1 serotype (**Figure 1E, Table S4**). These findings suggest that VP1 is the primary source of bovine CD8+ and CD4+ T cell epitopes, similar to the mouse system.

### Conserved T cell epitope presentation in BTV proteins by a laboratory murine model

Consistent with our analysis of multiple sequence alignments that revealed significant conservation in the VP1 and VP7 proteins of BTV, we identified highly conserved CD8+ and CD4+ T cell epitopes within these proteins, present across all 24 BTV serotypes (**Figure 2**)(**Table S2**). Based on previous research, we established a cut-off value of 62% to determine whether an epitope should be classified as conserved (Tengvall K 2019, Girdhar, Huang et al. 2022). Interestingly, all predicted CD8+ and CD4+ T cell epitopes in VP1 exhibited substantial conservation, with a minimum of 66.66% amino acid sequence similarity. Remarkably, VP1 contained 15 CD8+ T cell epitopes that were 100% conserved, with the remaining epitopes displaying conservation levels ranging from 66.66% to 88.88%. Similarly, VP1 harbored 13 CD4+ T cell epitopes that were 100% conserved, along with others exhibiting conservation levels ranging from 66.66% to 93.33% (**Figure 2A, 2B**). In contrast, the VP1 protein exhibited 13 CD4+ T cell epitopes that were 100% conserved, followed by 9 epitopes with 93.33%, 18 with 86.66%, 1 with 73.33%, and 5 with 66.66% conservation (**Figure 2C, 2D**). Regarding the VP7 protein, 2 CD8+ T cell epitopes were found to be 100% conserved, with 4 epitopes exhibiting 88.88% conservation, 3 with 77.77%, and 3 with 66.66%. Additionally, two CD8+ T cell epitopes displayed conservation levels below 50%, with conservation rates of 33.33% and 22.22%. Although the VP7 protein contained a higher frequency of CD8+ T cell epitopes, it contained only 1 CD4+ T cell epitope with 66.66% conservation. Consistent with expectations, no conserved CD8+ and CD4+ T cell epitopes were detected in the VP2 protein. Despite amino acid variability in the VP5 protein, one CD8+ T cell epitope with 66.66% conservation and 2 CD4+ T cell epitopes with 60% conservation were identified in this protein.

**Figure 2.**
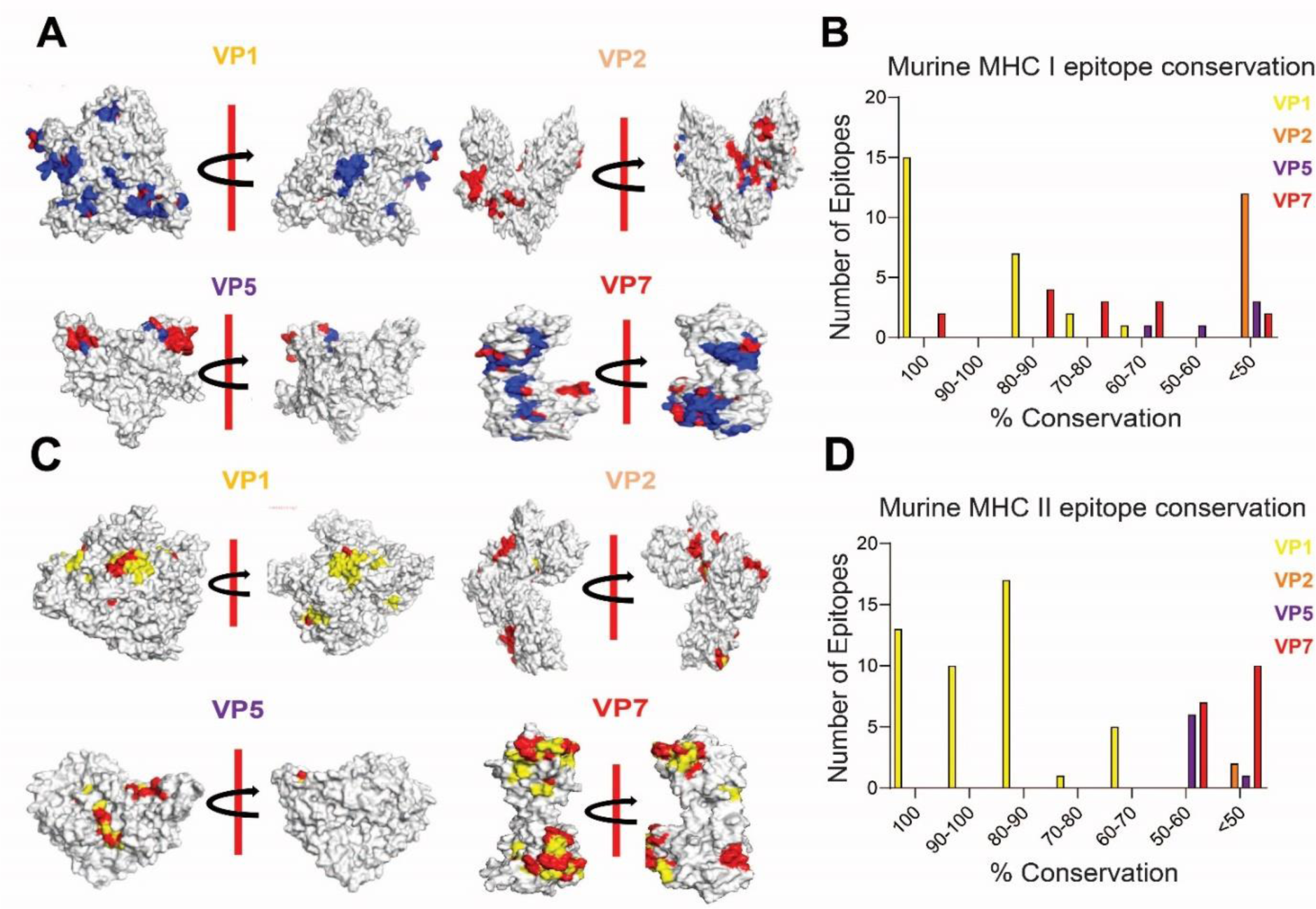
The presence of conserved CD8+ and CD4+ T cell epitopes corresponding to mouse system. **A.** Representation of conserved (highlighed in blue) and varied (highlighted in red) amino acid residues in the mouse CD8+ T cell epitopes of BTV structural proteins. The protein structures were predicted using Robetta server and both front and back views are presented at 180° rotation. **B.** Bar graph presenting the degree of amino acid conservation among MHC class I-restricted CD8+ T cell epitopes in BTV structural proteins. **C.** Representation of conserved (highlighed in blue) and varied (highlighted in red) amino acid residues in the mouse CD4+ T cell epitopes of BTV structural proteins. The protein structures were predicted using Robetta server and both front and back views are presented at 180° rotation. **D.** Bar graph presenting the degree of amino acid conservation among MHC class II-restricted CD4+ T cell epitopes in BTV structural proteins.

### Distribution of conserved CD8+ and CD4+ T cell epitopes in Bovines

Similar to the CD8+ and CD4+ T cell epitopes recognized by the murine system; we observed a comparable pattern of conserved epitopes in the bovine system (**Fig 3; Table S3)**. The VP1 protein exhibited 10 BoLA class I-specific CD8+ T cell epitopes that were 100% conserved, with an additional 10 epitopes displaying 88.88% conservation, while 6 epitopes exhibited 77.77% conservation and another 6 had 66.66% conservation (**Fig 3A, 3B)**. Only 1 epitope showed 55.55% conservation among all the predicted CD8+ T cell epitopes. Similarly, 11 CD4+ T cell epitopes were 100% conserved in the VP1 protein, followed by 9 epitopes with 93.33% conservation, 7 with 86.66%, 6 with 80%, and 11 with 73.33% (**Fig 3A, 3B).** A similar trend of CD8+ T cell epitope conservation was observed in the VP7 protein, which included 1 epitope each with 100%, 88.88%, and 55.55%, and 2 epitopes each with 77.77% and 66.66%. The VP7 protein also contained 1 CD4+ T cell epitope that was 100% conserved, along with 2 epitopes with 93.33%, 1 with 73.33%, and 2 with 66.66% conservation, covering all 24 BTV serotypes (**Fig 3C, 3D)**. As expected, no conserved CD8+ and CD4+ T cell epitopes were found in VP2. Surprisingly, we identified 4 CD8+ T cell epitopes, 1 with 88.88% and 3 epitopes with 66.66% conservation in the VP5 protein, the second most variable structural protein after VP2. In contrast, there were few CD4+ T cell epitopes in the protein, which were highly conserved to a certain extent (1 with 73.33%, 1 with 66.66%, and 1 with 60%)(**Fig 3D;Table S4)**.

**Figure 3.**
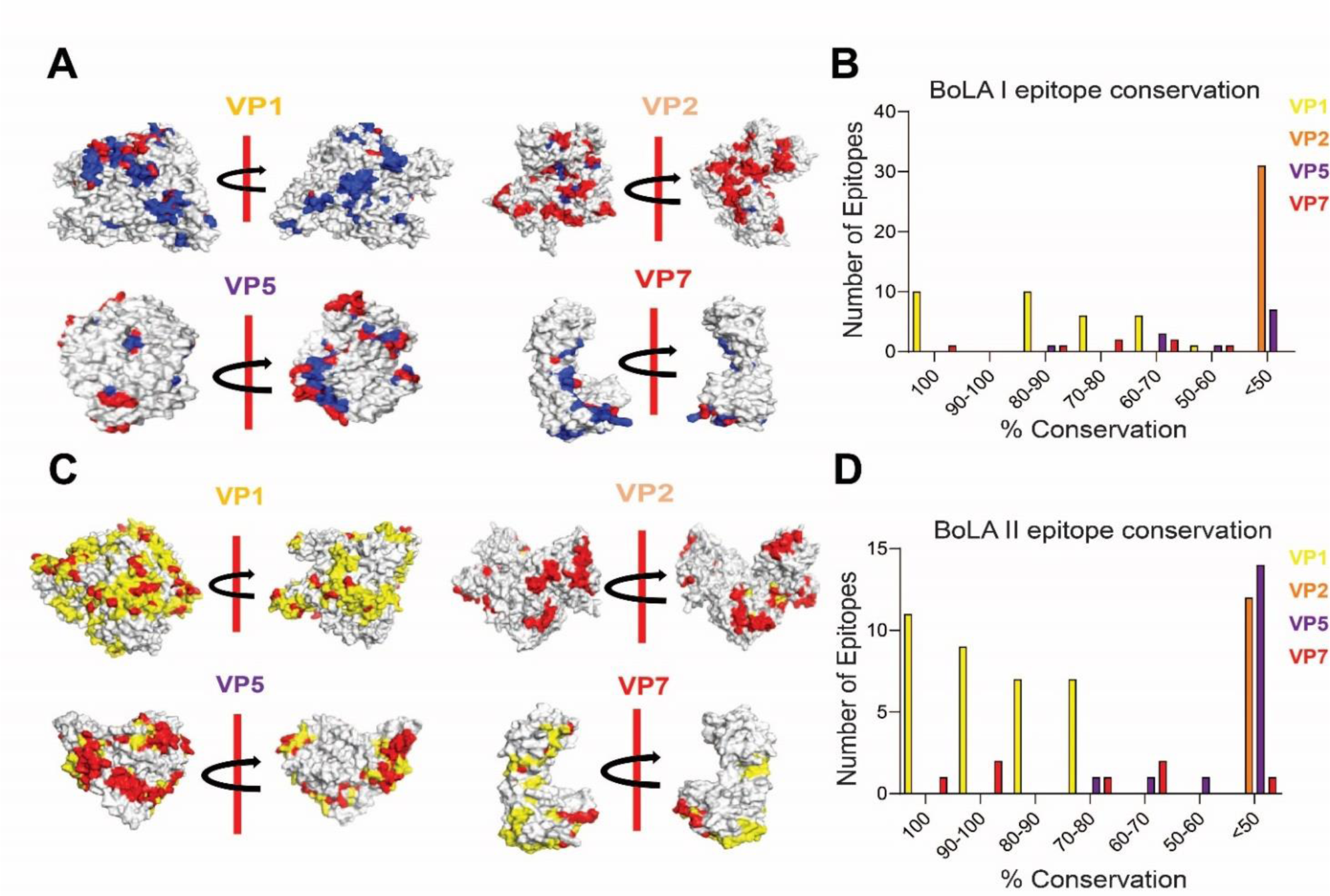
Presence of conserved CD8+ and CD4+ T cell epitopes corresponding to Bovine system. **A.** Representation of conserved (highlighed in blue) and varied (highlighted in red) amino acid residues in the bovine CD8+ T cell epitopes of BTV structural proteins. The protein structures were predicted using Robetta server and both front and back views are presented at 180° rotation. **B.** Bar graph presenting the degree of amino acid conservation among BoLA class I-restricted CD8+ T cell epitopes in BTV structural proteins. **C.** Representation of conserved (highlighed in blue) and varied (highlighted in red) amino acid residues in the BoLA CD4+ T cell epitopes of BTV structural proteins. The protein structures were predicted using Robetta server and both front and back views are presented at 180° rotation. **D.** Bar graph presenting the degree of amino acid conservation among BoLA class II-restricted CD4+ T cell epitopes in BTV structural proteins.

### VP1 and VP7 structural proteins are predominant source of conserved MHC- and BoLA-specific T cell epitopes

We observed a higher frequency of conserved CD8+ and CD4+ T cell epitopes in both murine and bovine systems within the VP1 and VP7 proteins (**Fig 4A**). Conversely, the VP2 and VP5 proteins contain more epitopes with less than 60% amino acid conservation. Interestingly, this pattern was consistent across both mouse and bovine systems, indicating that the VP1 and VP7 proteins are potential targets for consideration in BT vaccine development.

**Figure 4.**
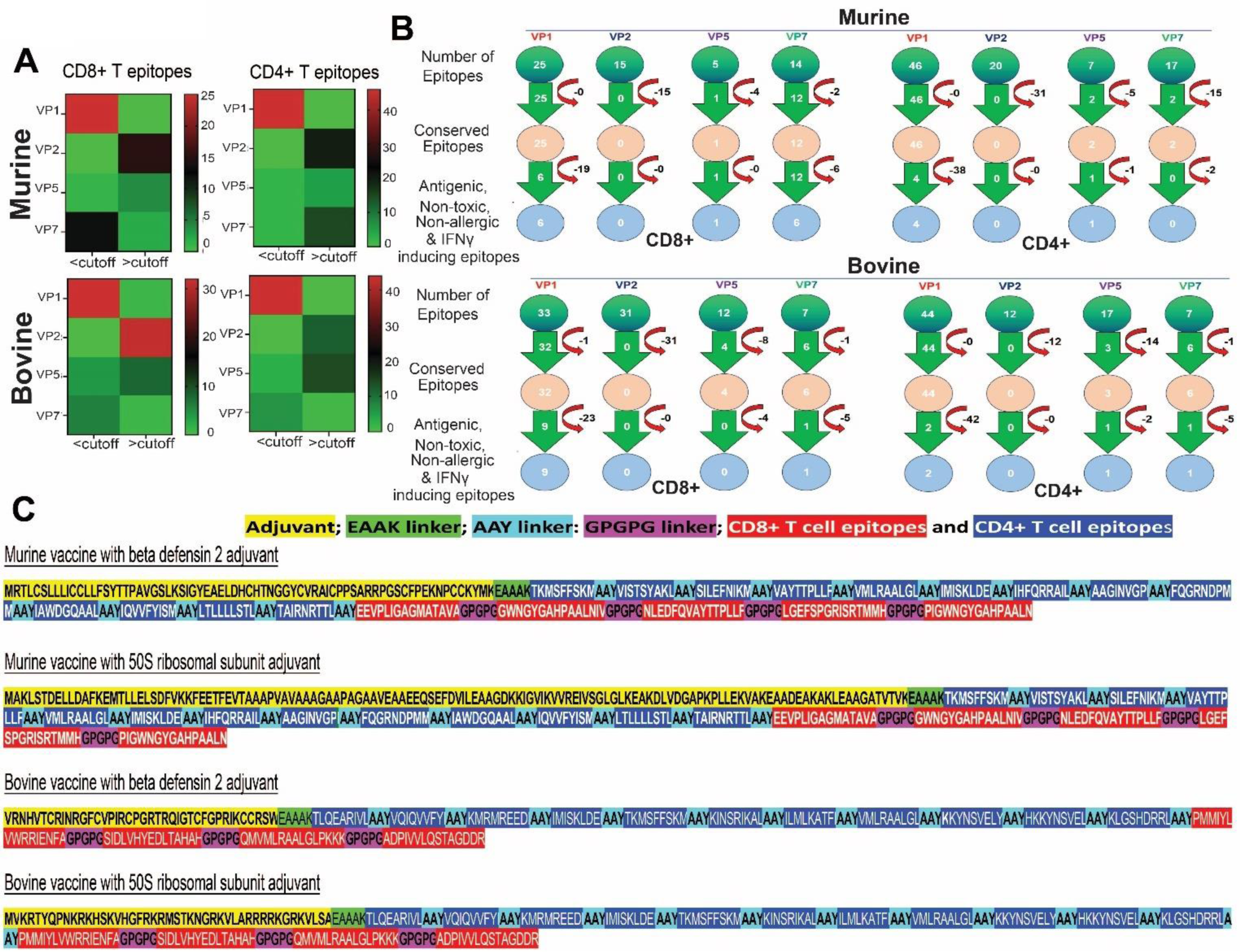
Conserved Epitopes among the structural proteins of BTV and the deisgn of a broad-spectrum BTV multi-epitope Vaccine. **A.** The VP1 and VP7 are the hotspots for conserved epitopes in both murine and bovine Systems. The heat maps show a predominance of CD4+ and CD8+ T cell epitopes, particularly within the VP1 protein. **B.** Screening of T cell epitopes for the design of a broad-spectrum BTV vaccine. **C.** Molecular design of a broad-spectrum BTV multi-epitope vaccine corresponding to murine and bovine systems. The CD8+ and CD4+ T cell epitopes were joined together with AAY and GPGPG linkers, respectively, whereas adjuvants are joined at the N-terminal of the vaccine construct using EAAAK linker.

### Selection of conserved epitopes and design of pan-BTV vaccine

After identifying highly conserved CD8+ and CD4+ T cell epitopes in both mouse and bovine systems, we subjected these epitopes to rigorous screening based on antigenicity, allergenicity, toxicity, and their ability to induce IFN-*γ*. This process yielded a total of 13 mouse CD8+ T cell epitopes—6 from VP1, 1 from VP5, and 6 from VP7 proteins—that met the criteria of being antigenic, non-allergenic, non-toxic, and IFN-*γ* inducers (**Fig 4B**). Similarly, we identified 5 CD4+ T cell epitopes—4 in VP1 and 1 in VP5 protein—that satisfied these criteria (**Fig 4B**). For the design of the bovine vaccine, we obtained a total of 11 CD8+ T cell epitopes—9 from VP1 and 2 from VP7 proteins (**Fig 4B**) —as well as 4 CD4+ T cell epitopes, 2 from VP1, and 1 each from VP5 and VP7 proteins (**Fig 4B**). Despite the final CD4+ T cell epitope from the VP5 protein not exhibiting high conservation (40 % conservation), we included it since it is conserved across many serotypes (all the serotypes with at least 40 % conservation), thereby increasing the overall number of CD4+ T cell epitopes in the bovine vaccine.

The selected epitopes, which demonstrated potent anti-viral T cell responses, were utilized to design *in silico* multi-epitope pan-BTV vaccines, as described in the Methods section (**Figure 4C**). Briefly, the conserved T cell epitopes were connected using suitable linkers, and TLR4 agonist adjuvants, such as beta-defensin 2 or 50s ribosomal protein, were integrated into the N-terminus region of the CD8+ CTL and CD4+ T helper polyepitopic region to boost immunogenicity (Pyasi, Sharma et al. 2021). With this approach, we developed 2 constructs for mice and 2 for bovines, each utilizing a distinct TLR4 agonist adjuvant: mouse vaccines with either beta-defensin 2 (**mVac-*β*-def**) or 50 S ribosomal subunit (**mVac-50sR**) adjuvants, and bovine vaccines with either beta-defensin 2 (**bVac-*β*-def**) or 50 S ribosomal protein (**bVac-50sR**) adjuvants.

### Physicochemical properties, Antigenicity, allergenicity, and solubility profiling of the vaccine constructs

The physicochemical properties of the vaccine constructs were analyzed, including molecular weights, theoretical pI, instability index, and GRAVY scores. The mVac-*β*-def, mVac-50sR, bVac-*β*-def, and bVac-50sR constructs have molecular weights of 34738.56, 40358.78, 28107.17, and 28892.16, respectively, with theoretical pI values of 8.86, 5.41, 9.79, and 10.25 (**Table 1**). Notably, all constructs exhibit stability, with instability index values of 33.67, 27.77, 36.24, and 44.65, respectively, although bVac-50sR appears slightly less stable compared to the others (**Table 1**). Additionally, GRAVY scores (measures the sum of hydropathy values of all amino acids in the protein) for all four constructs are 0.363, 0.353, 0.042, and −0.183, indicating the mouse vaccines are hydrophobic while bovine vaccines, bVac-*β*-def, and bVac-50sR, are neutral and hydrophilic in nature (**Table 1**). Antigenicity profiling reveals scores of 0.6385, 0.5678, 0.5736, and 0.5501, respectively, indicating high antigenicity for all vaccines (**Table 1**). Lastly, allergenicity profiling confirms that all vaccine constructs are non-allergenic (**Table 1**).

**Table 1.**
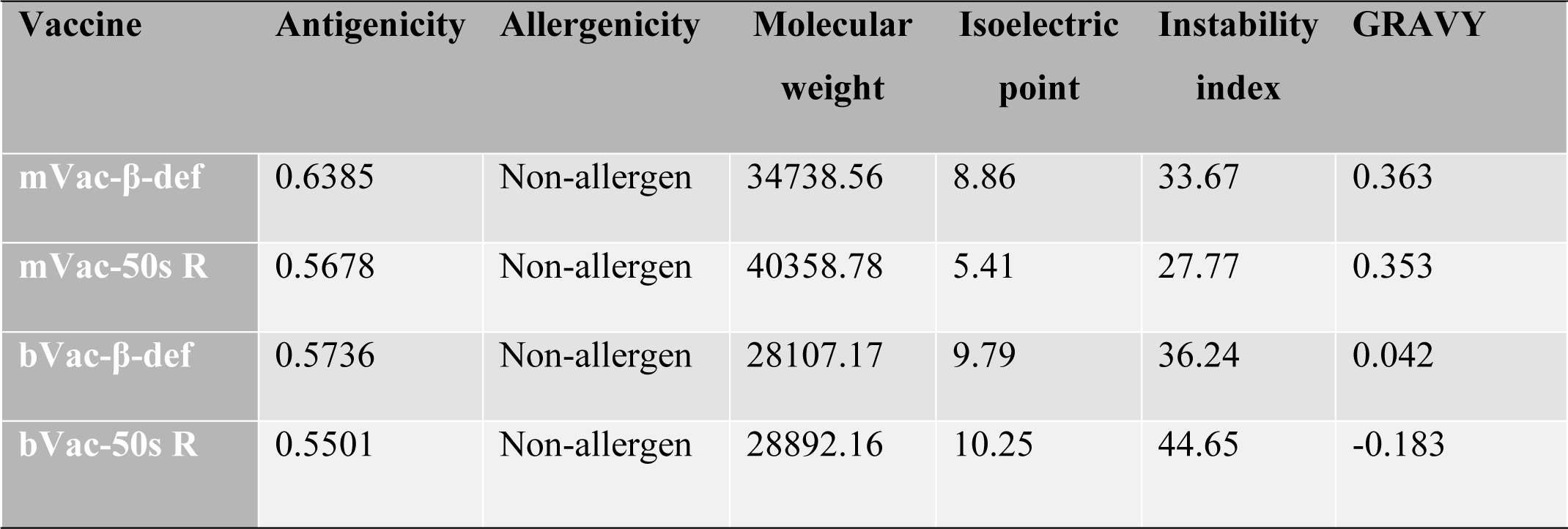
Immunological and physicochemical attributes of all the designed vaccine constructs.

### Structural modeling and evaluation of vaccine constructs

We utilized a protein modeling approach to generate the 3D structures of all four vaccine constructs for molecular docking studies (**Fig 5A**). These protein models underwent validation based on their phi (ϕ) and psi (Ψ) torsion angles through Ramachandran plot analysis. A protein model with over 90% of the residues in the most favorable regions is considered optimal. As depicted in **Fig 5B**, the Ramachandran plot analysis for mVac-*β*-def, mVac-50sR, bVac-*β*-def, and bVac-50sR-modeled structures reveals values within the favorable regions of 90.7%, 93.3%, 95.9%, and 97%, with additional values falling within the allowed regions of 5.9%, 5.5%, 3.6%, and 2.6%, respectively **(Table S5)**. Moreover, only 1.5%, 0.3%, 0%, and 0% were found in the generously allowed regions, and 1.9%, 0.9%, 0.5%, and 0.4% were scattered in disallowed regions of the Ramachandran plots. These findings confirm the excellent quality of the modeled 3D structures of the vaccine constructs, rendering them highly suitable for molecular docking studies.

**Figure 5.**
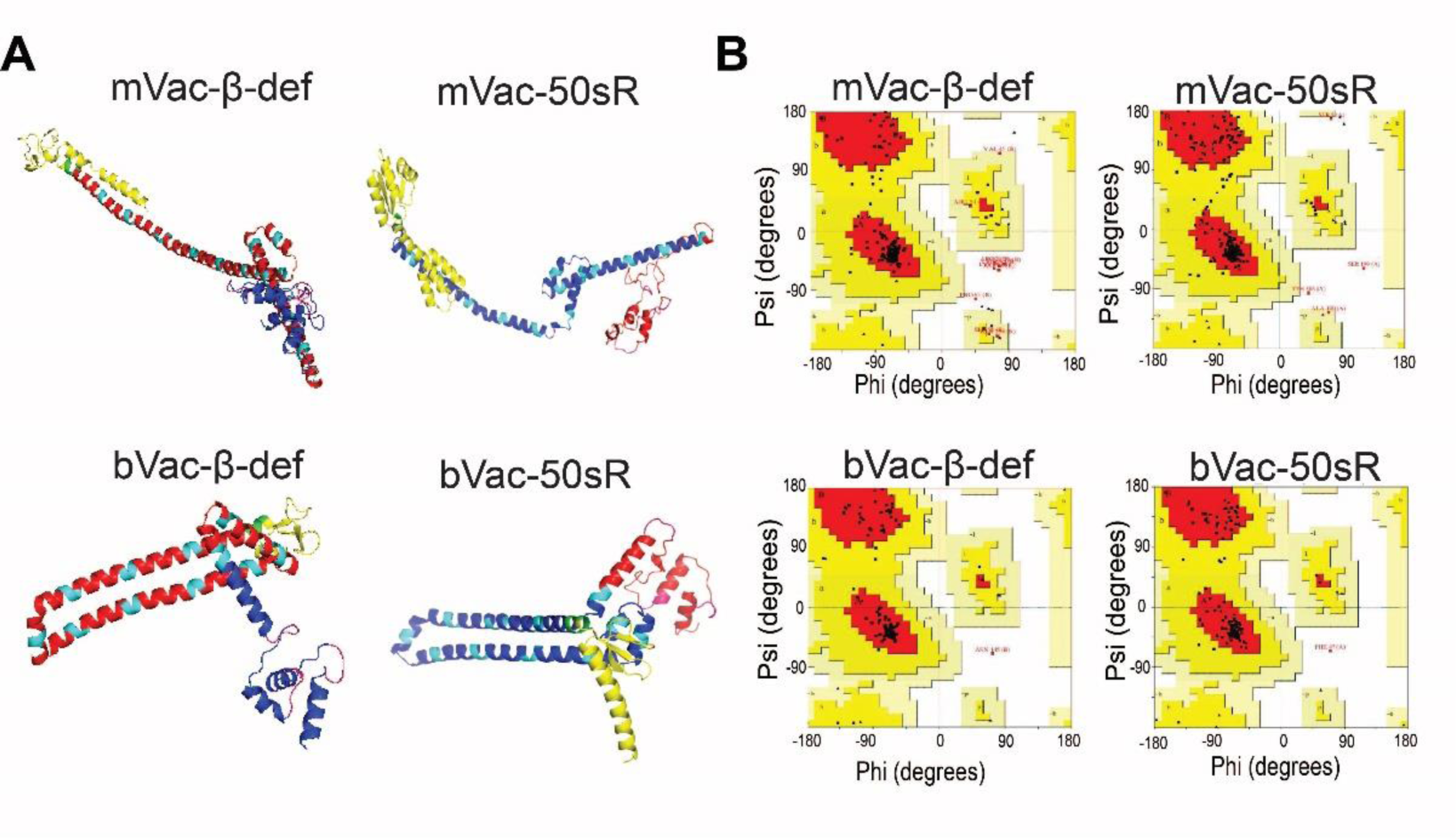
Protein modeling and evaluation of three-dimensional structures of the vaccine constructs. **A.** The three-dimensional structures of vaccine constructs obtained through protein modeling. **B.** Ramachandran plots showing the quality of modeled protein structures of vaccine constructs.

### Protein-Protein docking

To predict the molecular interaction patterns between all four vaccine constructs (mVac-*β*-def, mVac-50sR, bVac-*β*-def, and bVac-50sR) and their receptor, TLR-4, we conducted protein-protein docking using the ClusPro platform. The results revealed binding affinities of −-15.5 kcal/mol, −20.9 kcal/mol, - 10.9 kcal/mol, and −16.6 kcal/mol for mVac-*β*-def, mVac-50sR, bVac-*β*-def, and bVac-50sR, respectively. Notably, mVac-50sR exhibited the highest negative docking score and confidence score, indicating its superior interaction with the TLR4 molecule.

Further analysis of post-docking interactions revealed a series of molecular interactions between the vaccine constructs and TLR-4, encompassing hydrogen bonds, salt bridges, and non-bonded contacts (**Fig 6A, 6B**). As depicted in **Fig 6A**, the docking complex of TLR4 with mVac-*β*-def vaccine formed 16 hydrogen bonds, 01 salt bridge, and 226 non-bonded interactions. Conversely, the complex of TLR4 and mVac-50sR vaccine exhibited 05 hydrogen bonds and 280 non-bonded interactions. In the case of bVac-*β*-def vaccine and TLR4, 11 hydrogen bonds, 03 salt bridges, and 151 non-bonded contacts were observed (**Fig 6B**). Similarly, the docking complex of bVac-50sR vaccine and TLR4 formed 06 hydrogen bonds, 01 salt bridge, and 174 non-bonded contacts (**Fig 6B**). Notably, this docked complex had the highest number of hydrogen bonds (06), while the murine mVac-50sR and TLR4 complex exhibited the highest number of non-bonded interactions (280).

**Figure 6.**
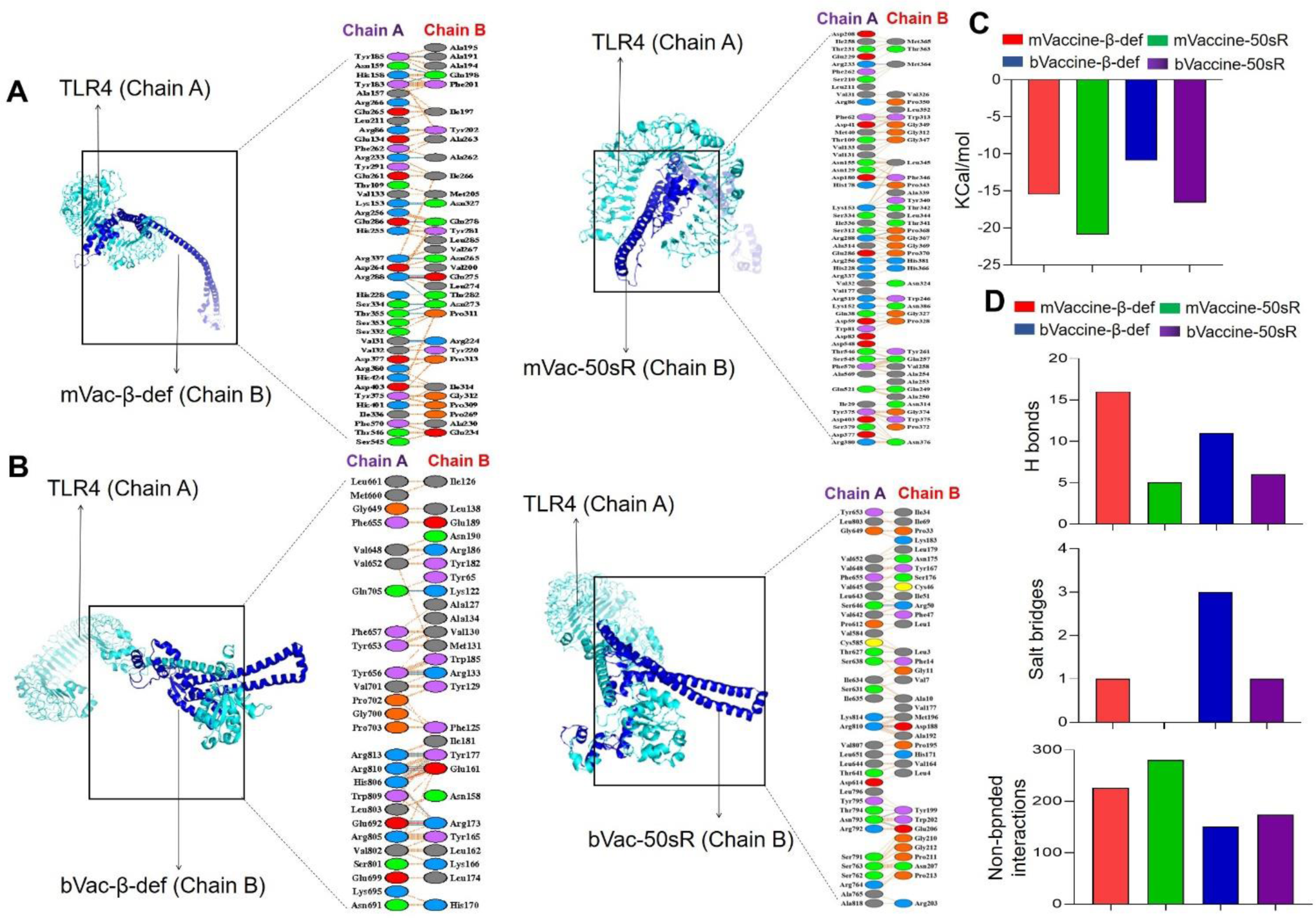
Molecular interactions between the vaccine constructs with mouse and bovine TLR4 structures. **A.** Visualization of molecular interactions between the mouse vaccine constructs (Chain B) in complex with TLR4 (Chain A) are represented in both 3-D and 2-D views. **B.** Visualization of molecular interactions between the bovine vaccine constructs (Chain B) in complex with TLR4 (Chain A) are represented in both 3-D and 2-D views. **C.** The Bar graph representing the Gibbs free energy values for the murine and bovine vaccine docked complexes. **D.** Bar graph representing the total number of hydrogen bonds, slat bridges and non-bonded interactions in the docked complex between the vaccine constructs and TLR4.

### Molecular dynamics simulation of docked complexes

MD simulations are one of the well-established methods for obtaining dynamic data at atomic spatial resolution (Gajula et al., 2016). In this study, MD simulations were performed to determine the structural stability of the docking complexes over 100 ns. To analyze the stability of the vaccine constructs in the binding regions of TLRs, root-mean-square deviation (RMSD) and root-mean-square fluctuation (RMSF) plots were calculated using several modules embedded in the Gromacs package. Here, we used representative mVac-50sR and bVac-50sR vaccines to assess their stability. As shown below, overall, the calculated RMSF plots corroborate the findings of the RMSD and docking analyses, indicating that the vaccine constructs interact significantly with their respective TLRs.

RMSD is a widely used method for measuring the stability of simulated systems and is valuable for assessing conformational changes within macromolecular backbone structures during MD simulations on a time scale (Sargsyan et al., 2017). To assess the stability of mVac-50sR and bVac-50sR vaccine constructs binding to their TLR-4, backbone RMSDs were analyzed graphically. As **Figure 7A** illustrates, both the docked complexes maintained consistent stability throughout the simulation, with RMSD values ranging from ∼0.95 to 4.1 nm. The average RMSD values for the mVac-50sR-TLR4 and bVac-50sR-TLR4 docking complexes were 1.72 and 1.83 nm, respectively. These complexes showed small fluctuations between ∼10 to 60 ns on ∼0.90 nm, suggesting relatively stable trends with minor conformational changes throughout the 100 ns simulation. Therefore, based on the minimal fluctuations observed and the low differences in computed RMSD scores, it can be inferred that the vaccine-TLR complexes remained stable.

**Figure 7.**
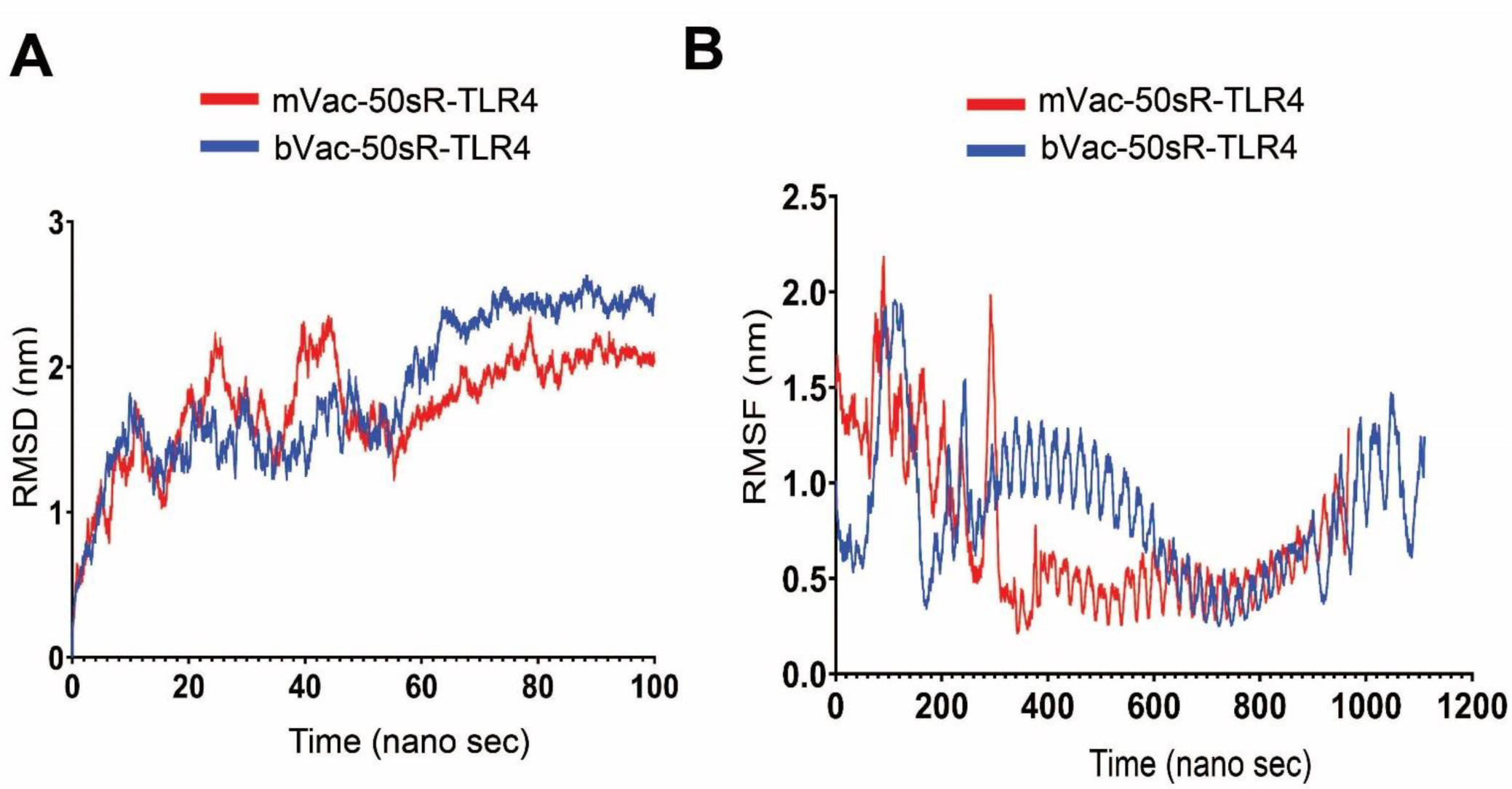
Molecular dynamic simulations of the vaccine-TLR4 docked complexes: **A.** The graph showing RMSD in the mouse and bovine vaccines and TLR4 docked complexes. **B.** The graph representing RMSF of the mouse and bovine vaccine and TLR4 docked complexes.

The RMSF plot analysis was conducted to assess the flexibility of individual residues within the simulated systems over time. As a rule, higher fluctuation scores indicate greater flexibility and less stable bonds, while lower values suggest well-structured segments in the protein-protein docking complexes (Gajula MNV KA 2016). **Figure 7B** depicts the calculated RMSF plots for the alpha-carbon atoms of all four simulated systems. As evident from this figure, both complexes, bVac-TLR4 (blue) and mVac-TLR4 (red) demonstrated the close pattern of the peaks between residues 1 to 300 on ∼2.2 nm. After 300 resides, no significant peak has been observed in these complexes. The average RMSF values for the mVac-50sR-TLR4 and bVac-50sR-TLR4 complexes were 0.73 and 0.85 nm, respectively. Both the simulated systems experienced relatively minor conformational changes throughout the simulations.

## Discussion

Vaccination is an effective measure to control the spread of BT infection among livestock (Feenstra and van Rijn 2017, van Rijn 2019, Rojas 2021). However, the efficacy of current live attenuated and inactivated vaccines is hampered by their serotype-specific nature, rendering them less effective against the evolving spectrum of BTV strains (Ranjan, Prasad et al. 2019). For instance, A decade ago, there were 24 classical BTV serotypes, but eight new ones have emerged since then. Consequently, the primary objective of BTV vaccination lies in eliciting a broad-spectrum immune response capable of combating all BTV serotypes while minimizing adverse reactions in hosts. In pursuit of this goal, our prior research successfully explored and developed pan-BTV vaccines targeting conserved epitopes within BTV nonstructural proteins, specifically NS1 and NS2 (Harish Babu Kolla 2023). Building upon this foundation, we sought to extend our efforts to encompass the structural proteins of BTV. Through rigorous epitope selection criteria, we identified highly conserved CD8+ and CD4+ T cell epitopes within VP1, VP5, and VP7 proteins of BTV, culminating in the development of in silico-based pan-BTV vaccines capable of conferring protection against all 24 classical serotypes. The exclusion of VP3 and VP4 proteins from our vaccine design was guided by their inhibitory nature, which could potentially impede the establishment of robust cross-reactive cellular responses (Chauveau, Doceul et al. 2013). Additionally, the structural proteins of BTV serotypes 25 to 32 were not considered due to incomplete sequence data available on PubMed. Nonetheless, given the inclusion of epitopes present across all 24 serotypes in our vaccine design, it is plausible that they exhibit conservation within the remaining BTV serotypes 25 to 32.

The integration of computational tools and immunoinformatics not only aids in identifying critical epitopes but also lays a robust foundation for advancing pan-BTV vaccine development. With meticulous scrutiny, we filtered conserved T cell epitopes based on antigenicity, allergenicity, toxicity, and IFN-*γ*-inducing abilities to pinpoint potential candidates for pan-BTV vaccine design. Utilizing these epitopes, we designed two vaccines each for the mouse (mVac-*β*-def with beta-defensin 2 and mVac-50sR with 50s ribosomal subunit adjuvant) and two for bovine system (bVac-*β*-def with beta-defensin 2 and bVac-50sR with 50s ribosomal subunit adjuvant), subjecting them to further analysis. All four designed vaccine constructs demonstrated an acceptable range of physicochemical properties at the protein sequence level, making them suitable for structural-based evaluation and annotations (**Table 1**). Based on the immunological profiling, all four vaccine constructs have been classified as probable antigens with high antigenic scores and non-allergens or not inducers of autoimmunity in the host. Furthermore, all the four vaccines are used for structural analysis using standard bioinformatics methods such as protein modeling, molecular docking, and molecular dynamic simulations. The most popular approach for homology modeling, also known as comparative modeling, was applied to predict the three-dimensional (3D) structural models of the vaccine constructs. Ramachandran plot analysis of the structural models is considered one of the most important approaches in recent vaccine studies (Samad, Meghla et al. 2022, Aiman, Ahmad et al. 2023). Based on their Ramachandran plot analysis all the four vaccine models were found to be stable with high quality and suitable for molecular docking studies.

The protein-protein docking is a crucial method for predicting molecular interactions between the TLRs and vaccine constructs (Vakser 2014). We targeted the TLR4 because of its potential role in skewing anti-viral immune response (Suh, Zhao et al. 2009). The protein-protein docking, and interaction analysis demonstrated that the designed vaccine constructs exhibit a strong molecular interaction pattern with their respective receptor molecules at the structural level. RMSD is a widely used method for measuring the stability of simulated systems and is valuable for assessing conformational changes within macro molecular backbone structures during MD simulations on a time scale (Sargsyan 2017). Based on the minimal fluctuations observed and the low differences in computed RMSD scores depicted in the RMSD plots, it can be inferred that the vaccine-TLR complexes remained stable. The RMSF plot analysis was conducted to assess the flexibility of individual residues within the simulated systems over time. As a rule, higher fluctuation scores indicate greater flexibility and less stable bonds, while lower values suggest well-structured segments in the protein-protein docking complexes (Gajula MNV KA 2016). In summary, the calculated RMSF plots corroborate the findings of the RMSD and docking analyses, indicating that the vaccine constructs interact significantly with their respective TLRs.

Collectively, the comprehensive profiling of our vaccine constructs underscores their potential as promising candidates for effective BT vaccine development. With a focus on cell-mediated immunity, which has demonstrated its importance in providing cross-protection against various BTV serotypes in prior research (Umeshappa, Singh et al. 2010, singh 2011, Umeshappa, Singh et al. 2011, Rojas, Rodriguez-Calvo et al. 2017, Potter 2019, Rijn 2019, Evseev and Magor 2021, Rodríguez-Martín D 2021, Wang, Tang et al. 2022, Zheng, Guo et al. 2022), our vaccine design strategy emphasizes targeting conserved immunogenic cytotoxic T lymphocyte (CTL) and T-helper-1 cell epitopes to develop a highly effective pan-BTV vaccine. Including T-helper epitopes is crucial, as CD4+ T-helper-1 cells play a vital role in supporting the function of anti-viral CD8+ CTLs (Umeshappa, Huang et al. 2009, Umeshappa and Xiang 2011, Sokke Umeshappa, Hebbandi Nanjundappa et al. 2012, Umeshappa, Xie et al. 2013). Upon vaccination, antigen-presenting cells (APCs) present epitopes for CD4+ T-helper-1 cell activation, subsequently activating APCs and providing helper signals for CD8+ CTL effector and memory responses. Thus, our vaccine is strategically designed to elicit robust cross-reactive cellular immunity against multiple BTV serotypes.

## Conclusion

The presence of multiple BTV serotypes challenges the effectiveness of existing vaccines, which predominantly target specific serotypes and may raise safety concerns. Drawing insights from broad-spectrum vaccine development for influenza, we devised and evaluated pan-BTV multi-epitope vaccines, aiming to cover all serotypes by targeting key structural proteins. These vaccines, incorporating conserved peptide regions, hold promise for inducing cross-protection against all serotypes, aiding global BTV control. Our comprehensive profiling underscores their potential as effective candidates, prioritizing cell-mediated immunity and focusing on conserved immunogenic epitopes. This approach, supported by prior research, emphasizes the importance of harnessing CD4+ T-helper-1 cells and CD8+ CTLs for optimal vaccine efficacy, thereby contributing to ongoing BT vaccine research and improving BTV immunoprophylaxis.

## Acknowledgement

The current study is pursued in collaboration with the Centre for Animal Disease Research and Diagnosis, Indian Veterinary Research Institute, Uttar Pradesh, India. The authors are grateful for start-up funding from the Faculty of Medicine, Dalhousie University. CSU is supported by the prestigious Canada Research Chair Tier 2 Award from Canadian Institutes of Health Research. HBK is supported from 2022 DMRF I3V Graduate Studentship. PPCM acknowledges support from EU H20:20 grant ‘PALE-Blu’ (project number 727393-2).

## Authors contributions

H.B.K generated the data in figures 1, 2, 3, 4, 5 and 6, and supplementary table 1-4 and 7 with contributions from C.S.U and A.K. A.K generated the data in figure 7, and supplementary tables 5-6 with contributions from M.D, and H.B.K under the supervision of D.K and C.S.U. K.P.S, R.H.N and P.P.C.M contributed to the data interpretation, figures preparation, and manuscript writing. C.S.U. designed the study, supervised, and coordinated its execution, and wrote the manuscript along with H.B.K and A.K.

