## Supplementary tables for "Unleashing the Immune Arsenal: Development of Broad-spectrum Multiepitope Bluetongue Vaccine Targeting Conserved T Cell Epitopes of Structural Proteins"

**Supplementary Table 1.** MHC class I specific CD8+ T cell epitopes in the BTV1 structural proteins.

| Allele | Peptide | ic50 | Percentile | % Conservation |
|---|---|---|---|---|
| **VP2** | | | | |
| H-2-Kb | FNKWIIAPM | 496.72 | 0.57 | 11.11 |
| H-2-Kb | IAYLEYMVF | 48.23 | 0.09 | 0 |
| H-2-Kb | ILLKFSGHV | 201.43 | 0.27 | 33.33 |
| H-2-Kb | IMYLNFLPL | 6.84 | 0.02 | 28.57 |
| H-2-Db | | 24.69 | 0.03 | |
| H-2-Kb | INFGRGQKV | 287.47 | 0.35 | 0 |
| H-2-Kb | LNFLPLYFL | 135.11 | 0.2 | 22.22 |
| H-2-Kb | LSIRFQEAI | 228.25 | 0.29 | 33.33 |
| H-2-Db | LSPITADPI | 141.42 | 0.07 | 0 |
| H-2-Kb | MMWNHLVRI | 93.99 | 0.15 | 0 |
| H-2-Kb | RFYDIRPAL | 286.49 | 0.35 | 11.11 |
| H-2-Kb | RTALWYNPI | 323.93 | 0.4 | 0 |
| H-2-Kb | SHRQWSIPL | 494.34 | 0.56 | 22.22 |
| H-2-Kb | TMPEYFNKW | 461.83 | 0.53 | 22.22 |
| H-2-Kb | VIAEFFPTY | 218.09 | 0.28 | 11.11 |
| H-2-Kb | YIYGRVNLF | 245.21 | 0.3 | 33.33 |
| **VP5** | | | | |
| H-2-Kb | EAYREFLNL | 45.54 | 0.08 | 44.44 |
| H-2-Kb | IHFQRRAIL | 90.2 | 0.15 | 66.66 |
| H-2-Kb | VHPIYLGSL | 170.83 | 0.24 | 55.55 |
| H-2-Kb | ISKAFGTQM | 218.11 | 0.28 | 0 |
| H-2-Kb | ILPRFKKAM | 377.89 | 0.46 | 22.22 |
| **VP7** | | | | |
| H-2-Db | AAGINVGPI | 30.04 | 0.03 | 88.88 |
| H-2-Kb | AAIARAAYV | 444.25 | 0.52 | 22.22 |
| H-2-Db | FAMHGVNPM | 41.44 | 0.03 | 33.33 |
| H-2-Db | FQGRNDPMM | 476.48 | 0.13 | 100 |
| H-2-Kb | IAINRYNGL | 13.07 | 0.03 | 100 |
| H-2-Kb | IAWDGQAAL | 320.38 | 0.39 | 66.66 |
| H-2-Kb | IQVVFYISM | 34.24 | 0.07 | 88.88 |
| H-2-Kb | ISPDYTQHM | 83.04 | 0.14 | 88.88 |
| H-2-Kb | KTLNQYPAL | 97.92 | 0.16 | 88.88 |
| H-2-Kb | LTLLLLSTL | 426.67 | 0.5 | 66.66 |
| H-2-Kb | MIYLVWRRI | 130.09 | 0.2 | 77.77 |
| H-2-Kb | RNEMFFMCL | 251.82 | 0.31 | 77.77 |
| H-2-Db | TAIRNRTTL | 7.36 | 0.01 | 77.77 |
| H-2-Kb | TVMRACATL | 233.81 | 0.29 | 66.66 |

**Supplementary Table 2.** MHC class II specific CD4+ T cell epitopes in the BTV1 structural proteins.

| Allele | Peptide Sequence | IC50 | Percentile Rank | % Conservation |
|---|---|---|---|---|
| **VP2** | | | | |
| H2-IAb | **EYMVFFPSKAIRLSK** | 133.12 | 0.28 | 0 |
| H2-IAb | **YMVFFPSKAIRLSKL** | 134.60 | 0.29 | 0 |
| H2-IAb | **LEYMVFFPSKAIRLS** | 164.16 | 0.44 | 0 |
| H2-IAb | **YLEYMVFFPSKAIRL** | 232.16 | 0.73 | 0 |
| H2-IAb | **MVFFPSKAIRLSKLN** | 251.06 | 0.86 | 0 |
| H2-IAb | **VFFPSKAIRLSKLNE** | 479.48 | 2.20 | 0 |
| H2-IAb | **QEEYIYGRV**NLF**DFV** | 550.82 | 2.60 | 20 |
| H2-IAb | **EQEEYIYGRV**NLF**DF** | 652.44 | 3 | 20 |
| H2-IAb | **LDGIVWYL**P**ITH**P**NK** | 733.09 | 3.40 | 13.33 |
| H2-IAb | **GIVWYL**P**ITH**P**NKCI** | 771.48 | 3.50 | 13.33 |
| H2-IAb | **REQEEYIYGRV**NLF**D** | 819.17 | 3.80 | 20 |
| H2-IAb | **FNKWIIA**P**MFNANVR** | 862.69 | 4.10 | 6.66 |
| H2-IAb | **EYFNKWIIA**P**MFNAN** | 878.70 | 4.20 | 6.66 |
| H2-IAb | **DGIVWYL**P**ITH**P**NKC** | 887.36 | 4.20 | 13.33 |
| H2-IAb | **EEYIYGRV**NLF**DFVA** | 888.02 | 4.20 | 20 |
| H2-IAb | **NKWIIA**P**MFNANVRI** | 891.32 | 4.20 | 6.66 |
| H2-IAb | **YFNKWIIA**P**MFNANV** | 906.44 | 4.30 | 13.33 |
| H2-IAb | **GRIRLRFPLSARHLK** | 921.80 | 4.30 | 0 |
| H2-IAb | **IVWYL**P**ITH**P**NKCIV** | 947.54 | 4.50 | 13.33 |
| H2-IAb | **FLDGIVWYL**P**ITH**PN | 950.15 | 4.50 | 13.33 |
| **VP5** | | | | |
| H2-IAb | EVP**LI**G**A**G**M**AT**AV**AT | 460.03 | 2.10 | 60 |
| H2-IAb | VP**LI**G**A**G**M**AT**AV**AT**G** | 552.13 | 2.60 | 53.33 |
| H2-IAb | EEVP**LI**G**A**G**M**AT**AV**A | 552.55 | 2.60 | 60 |
| H2-IAb | P**LI**G**A**G**M**AT**AV**AT**G**R | 672.63 | 3.10 | 53.33 |
| H2-IAb | **LI**G**A**G**M**AT**AV**AT**G**RA | 731.67 | 3.40 | 53.33 |
| H2-IAb | **MMK**F**KI**P**RA**Q**Q**P**QI**H | 775.56 | 3.50 | 33.33 |
| H2-IAb | **S**EEVP**LI**G**A**G**M**AT**AV** | 852.53 | 4 | 53.33 |
| **VP7** | | | | |
| H2-IAb | WFMR**A**AQ**AA**T**AVVC**G | 108.16 | 0.21 | 53.33 |
| H2-IAb | **MPWPL**T**A**AI**ARAAYV** | 132.97 | 0.28 | 20 |
| H2-IAb | P**MPWPL**T**A**AI**ARAAY** | 170.86 | 0.48 | 26.66 |
| H2-IAb | FMR**A**AQ**AA**T**AVVC**GP | 189.14 | 0.52 | 53.33 |
| H2-IAb | **N**P**MPWPL**T**A**AI**ARAA** | 270.87 | 0.96 | 26.66 |
| H2-IAb | **MDTIAARAL**TV**M**R**A**C | 457.85 | 2.10 | 26.66 |
| H2-IAb | MR**A**AQ**AA**T**AVVC**GPD | 580.79 | 2.70 | 60 |
| H2-IAb | RIE**N**F**AMAQ**GNS**QQ**T | 671.74 | 3.10 | 53.33 |
| H2-IAb | **R**PEF**AM**HG**VN**P**MPWP** | 681.15 | 3.10 | 40 |
| H2-IAb | IE**N**F**AMAQ**GNS**QQ**T**Q** | 689.70 | 3.20 | 46.66 |
| H2-IAb | RRIE**N**F**AMAQ**GNS**QQ** | 712.81 | 3.30 | 53.33 |

| | | | | |
|---|---|---|---|---|
| H2-IAb | L**R**PEF**AM**HG**VN**P**MPW** | 737.76 | 3.40 | 46.66 |
| H2-IAb | **DTIAARAL**TV**M**R**A**C**A** | 753.63 | 3.40 | 26.66 |
| H2-IAb | PEF**AM**HG**VN**P**MPWPL** | 842.60 | 3.90 | 40 |
| H2-IAb | VL**R**PEF**AM**HG**VN**P**MP** | 958.48 | 4.50 | 53.33 |
| H2-IAb | EF**AM**HG**VN**P**MPWPL**T | 987.88 | 4.60 | 40 |
| H2-IAb | WRRIE**N**F**AMAQ**GNS**Q** | 999.30 | 4.70 | 60 |

**Supplementary Table 3.** BoLA class I specific CD8+ T cell epitopes in the BTV1 structural proteins.

| Allele | Peptide | ic50 | Percentile Rank | % Conservation |
|---|---|---|---|---|
| **VP2** | | | | |
| BoLA-6:01301 | AQRQSDDPM | 41.47 | 0.15 | 22.22 |
| BoLA-1:02301 | | 208.36 | 0.05 | |
| BoLA-4:02401 | CLHTRTMMW | 323.91 | 0.02 | 22.22 |
| BoLA-6:01301 | EQFKMHKIL | 82.9 | 0.27 | 11.11 |
| BoLA-6:01301 | IMYLNFLPL | 23.16 | 0.09 | 22.22 |
| BoLA-1:02301 | | 221.02 | 0.06 | |
| BoLA-6:01302 | | 242.34 | 0.04 | |
| BoLA-1:02301 | KKAGYAEVL | 277.02 | 0.08 | 0 |
| BoLA-6:01301 | KLNEAHAKI | 259.2 | 0.66 | 0 |
| BoLA-6:01301 | KMVEGLTHL | 22.11 | 0.09 | 0 |
| BoLA-6:01301 | KQESIRTAL | 23.48 | 0.1 | 0 |
| BoLA-6:01301 | LLRGYEFTI | 475.52 | 1.0 | 11.11 |
| BoLA-6:01301 | LQRLTLARF | 185.19 | 0.5 | 0 |
| BoLA-1:02301 | | 432.87 | 0.13 | |
| BoLA-6:01301 | MMWNHLVRI | 66.18 | 0.24 | 0 |
| BoLA-1:02301 | | 477.71 | 0.16 | |
| BoLA-6:01301 | MVKRTLSPI | 176.98 | 0.48 | 11.11 |
| BoLA-6:01301 | REMRGKEKL | 201.49 | 0.52 | 0 |
| BoLA-6:01301 | RFYDIRPAL | 135.77 | 0.4 | 11.11 |
| BoLA-2:01201 | RGIVQIPKK | 320.87 | 0.03 | 0 |
| BoLA-6:01301 | RGKEKLNVI | 450.51 | 0.97 | 0 |
| BoLA-6:01301 | RLNHSTREI | 363.07 | 0.83 | 0 |
| BoLA-6:01301 | RQWSIPLLL | 8.3 | 0.03 | 22.22 |
| BoLA-1:02301 | | 67.84 | 0.02 | |
| BoLA-6:01302 | | 97.8 | 0.02 | |
| BoLA-1:02301 | SHRQWSIPL | 199.5 | 0.05 | 33.33 |
| BoLA-1:02301 | SKKADTMSY | 184.98 | 0.04 | 11.11 |
| BoLA-4:02401 | SMMRSWYDW | 70.78 | 0.01 | 11.11 |
| BoLA-6:01301 | SVRAGRIRL | 447.41 | 0.97 | 0 |
| BoLA-1:02301 | TKLGDVYSM | 476.67 | 0.16 | 33.33 |
| BoLA-6:01301 | TMSYHVEPI | 286.73 | 0.71 | 11.11 |
| BoLA-1:02301 | TRVWWSNPY | 459.52 | 0.15 | 33.33 |
| BoLA-6:01301 | VIRDDIASL | 267.44 | 0.68 | 11.11 |
| BoLA-6:01301 | VMRGKMPEV | 120.88 | 0.36 | 0 |
| BoLA-6:01301 | VQWMMKDSM | 52.63 | 0.19 | 0 |
| BoLA-1:02301 | | 275.16 | 0.08 | |
| BoLA-2:01201 | VTMPEYFNK | 392.03 | 0.03 | 22.22 |
| BoLA-6:01301 | YIYGRVNLF | 313.75 | 0.76 | 33.33 |
| BoLA-1:02301 | YKRGFPEHL | 448.53 | 0.14 | 0 |
| **VP5** | | | | |

| | | | | |
|---|---|---|---|---|
| BoLA-6:01301 | A**L**K**F**G**C**K**VL** | 494.32 | 1.2 | 44.44 |
| BoLA-1:02301 | **A**Q**Q**P**QI**H**V**Y | 157.2 | 0.04 | 44.44 |
| BoLA-6:01301 | E**QRN**E**LV**R**L** | 285.08 | 0.71 | 33.33 |
| BoLA-6:01301 | **IMRDRRQM**I | 66.47 | 0.24 | 11.11 |
| BoLA-6:01301 | K**L**K**K**VI**N**AL | 10.3 | 0.04 | 66.66 |
| BoLA-6:01302 | | 228.41 | 0.04 | |
| BoLA-6:01301 | **LM**H**IKNE**I**L** | 9.57 | 0.03 | 22.22 |
| BoLA-6:01302 | | 453.76 | 0.06 | |
| BoLA-4:02401 | **QI**H**V**Y**SA**PW | 484.52 | 0.03 | 44.44 |
| BoLA-6:01301 | RAIEGAYK**L** | 341.11 | 0.8 | 88.88 |
| BoLA-2:01201 | **R**SL**NR**FGKK | 173.61 | 0.02 | 66.66 |
| BoLA-1:02301 | S**KTV**HPI**Y**L | 446.31 | 0.14 | 55.55 |
| BoLA-6:01301 | **TQM**H**TR**R**LV** | 378.46 | 0.85 | 22.22 |
| BoLA-6:01301 | **V**QGSVHS**II** | 418.23 | 0.92 | 66.66 |
| **VP7** | | | | |
| BoLA-6:01301 | **A**Q**R**N**E**MFFM | 55.8 | 0.2 | 66.66 |
| BoLA-1:02301 | | 142.7 | 0.04 | |
| BoLA-6:01301 | IAINRYNGL | 446.93 | 0.97 | 100 |
| BoLA-6:01301 | KTLNQYP**A**L | 155.09 | 0.44 | 88.88 |
| BoLA-6:01301 | S**LA**Q**R**N**E**MF | 281.42 | 0.7 | 55.55 |
| BoLA-6:01301 | TLQ**EA**RIVL | 382.43 | 0.86 | 77.77 |
| BoLA-6:01301 | TV**M**R**A**C**A**TL | 264.95 | 0.67 | 66.66 |
| BoLA-6:01301 | **V**QIQVVFY**I** | 226.7 | 0.59 | 77.77 |

**Supplementary Table 4.** BoLA class II specific CD4+ T cell epitopes in the BTV1 structural proteins.

| | | | | | |
|---|---|---|---|---|---|
| **VP2** | | | | | |
| BoLA-DRB3_1201 | **DHELEIFGESIVDIS** | 1.000 | 0.684825 | 0.88 | 0 |
| BoLA-DRB3_1201 | **DVAYGQMINEMING**G | 1.000 | 0.889147 | 0.28 | 6.66 |
| BoLA-DRB3_0101 | K**PTYDIVVHAERRDR** | 0.620 | 0.649564 | 0.96 | 6.66 |
| BoLA-DRB3_1001 | **M**P**E**Y**FNKWIIA**P**MFN** | 1.000 | 0.733469 | 0.37 | 20 |
| BoLA-DRB3_1101 | **NKCIVAIEVS**D**ERVP** | 1.000 | 0.971167 | 0.02 | 6.66 |
| BoLA-DRB3_1601 | P**LYFLVGDNMI**Y**S**H**R** | 0.567 | 0.791218 | 0.45 | 20 |
| BoLA-DRB3_1201 | **REMLKY**Y**ANTTVYDG** | 0.993 | 0.852492 | 0.37 | 6.66 |
| BoLA-DRB3_1601 | S**IR**FQ**EAIDNK**F**RQH** | 1.000 | 0.846016 | 0.31 | 26.66 |
| BoLA-DRB3_1201 | **TMS**Y**HVEPIEDASKG** | 1.000 | 0.809814 | 0.47 | 6.66 |
| BoLA-DRB3_1601 | **VLTIDF**E**K**D**AKLTTN** | 0.993 | 0.946640 | 0.09 | 13.33 |
| BoLA-DRB3_0101 | **YDIVVHAERRDRSQP** | 0.993 | 0.868815 | 0.28 | 0 |
| BoLA-DRB3_0101 | **YSEGIVSHRVCKKNL** | 1.000 | 0.892155 | 0.23 | 0 |
| **VP5** | | | | | |
| BoLA-DRB3_2002 | **A**Y**R**EF**LNLAISKAFG** | 1.000 | 0.889590 | 0.38 | 20 |
| BoLA-DRB3_0101 | GK**V**I**R**SL**NR**FGKK**VG** | 0.973 | 0.652679 | 0.94 | 60 |
| BoLA-DRB3_0101 | **G**RAIEGAYK**L**K**K**VI**N** | 1.000 | 0.991217 | 0.01 | 73.33 |
| BoLA-DRB3_1201 | **IKEK**F**EKE**L**EEV**Y**N**F | 1.000 | 0.671822 | 0.93 | 26.66 |
| BoLA-DRB3_1601 | **K**NA**IEV**E**RDGMQE**EA | 1.000 | 0.799382 | 0.43 | 26.66 |
| BoLA-DRB3_1201 | L**EEV**Y**N**F**YNGEANAE** | 0.880 | 0.679128 | 0.90 | 20 |
| BoLA-DRB3_2002 | LK**K**VI**N**AL**S**GIDL**T**H | 1.000 | 0.823960 | 0.65 | 66.66 |
| BoLA-DRB3_1001 | **LT**E**A**Y**R**EF**LNLAISK** | 1.000 | 0.778537 | 0.29 | 26.66 |
| BoLA-DRB3_1201 | **NHK**E**LM**H**IKNE**I**L**P**R** | 1.000 | 0.868211 | 0.33 | 33.33 |
| BoLA-DRB3_1001 | NKA**VTS**Y**NKILTE**E**D** | 1.000 | 0.766740 | 0.31 | 26.66 |

| | | | | | |
|---|---|---|---|---|---|
| BoLA-DRB3_1101 | **P**D**NA**L**AVSV**L**I**K**ERA** | 1.000 | 0.716061 | 0.44 | 20 |
| BoLA-DRB3_1101 | **RD**KI**D**AL**KN**A**IEV**E**R** | 0.993 | 0.566900 | 0.88 | 46.66 |
| BoLA-DRB3_0101 | **R**P**SVVSTI**L**EYRAKE** | 0.987 | 0.720759 | 0.71 | 13.33 |
| BoLA-DRB3_1201 | **SIDL**V**HY**EDL**TA**H**A**H | 0.993 | 0.854740 | 0.36 | 40 |
| BoLA-DRB3_1201 | **T**E**A**Y**R**EF**LNLAISKA** | 1.000 | 0.970626 | 0.10 | 26.66 |
| BoLA-DRB3_1001 | **V**R**L**K**YNDKIKEK**F**EK** | 0.820 | 0.516307 | 0.88 | 20 |
| BoLA-DRB3_1601 | Y**N**F**YNGEANAEIEDE** | 1.000 | 0.822089 | 0.37 | 13.33 |
| **VP7** | | | | | |
| BoLA-DRB3_2002 | IAINRYNGLTLRGVT | 1.000 | 0.836355 | 0.60 | 100 |
| BoLA-DRB3_1501 | LAD**V**Y**T**VL**R**PEF**AMH** | 1.000 | 0.731772 | 0.68 | 66.66 |
| BoLA-DRB3_2002 | | 1.000 | 0.864128 | 0.48 | |
| BoLA-DRB3_1001 | PMMIY**LV**WRRIE**N**F**A** | 0.993 | 0.649253 | 0.53 | 73.33 |
| BoLA-DRB3_1601 | QVVFY**I**SMDKTLNQY | 1.000 | 0.911634 | 0.16 | 93.33 |
| BoLA-DRB3_0101 | RGVTMRP**T**S**LA**Q**R**N**E** | 1.000 | 0.709026 | 0.75 | 66.66 |
| BoLA-DRB3_1201 | **R**PEF**AM**HG**VN**P**MPWP** | 1.000 | 0.754583 | 0.64 | 46.66 |
| BoLA-DRB3_1501 | TIGVLATPE**I**PFTTE | 1.000 | 0.635758 | 1.00 | 93.33 |

**Supplementary Table 5.** List of finalized CD8+ and CD4+ T cell epitopes for the design of pan-BTV multi-epitope mouse vaccine.

| | Allele | Epitope | Antigenicty score | Antigenicty | Allergenicity | Toxicity | |
|---|---|---|---|---|---|---|---|
| **Mouse CD8+ T cell epitopes** | **VP1** | | | | | | |
| | **H-2-Kb** | TKMSFFSKM | **1.1372** | Antigen | Non-allergen | Non-Toxin | |
| | **H-2-Kb** | VISTSYAKL | **0.5864** | Antigen | Non-allergen | Non-Toxin | |
| | **H-2-Kb** | SILEFNIKM | **1.6253** | Antigen | Non-allergen | Non-Toxin | |
| | **H-2-Kb** | VAYTTPLLF | **0.7449** | Antigen | Non-allergen | Non-Toxin | |
| | **H-2-Kb** | VMLRAALGL | **0.6255** | Antigen | Non-allergen | Non-Toxin | |
| | **H-2-Db** | IMISKLDEI | **0.5623** | Antigen | Non-allergen | Non-Toxin | |
| | **VP5** | | | | | | |
| | **H-2-Kb** | IHFQRRAIL | 1.5373 | Antigen | Non-allergen | Non-Toxin | |
| | **VP7** | | | | | | |
| | **H-2-Db** | AAGINVGPI | 1.5614 | Antigen | Non-allergen | Non-Toxin | |
| | **H-2-Db** | FQGRNDPMM | 1.2268 | Antigen | Non-allergen | Non-Toxin | |
| | **H-2-Kb** | IAWDGQAAL | 0.7831 | Antigen | Non-allergen | Non-Toxin | |
| | **H-2-Kb** | IQVVFYISM | 0.8668 | Antigen | Non-allergen | Non-Toxin | |
| | **H-2-Kb** | LTLLLLSTL | 0.6520 | Antigen | Non-allergen | Non-Toxin | |
| | **H-2-Db** | TAIRNRTTL | 0.8312 | Antigen | Non-allergen | Non-Toxin | |
| **Mouse CD4+ T cell epitopes** | **VP1** | | | | | | |
| | **Allele** | **Epitope** | **Antigenicity score** | **Antigenicity** | **Allergenicity** | **Toxicity** | **IFNg inducing** |
| | **H2-IAb** | GWNGYGAHPAALNIV | 0.7697 | Antigen | Non-allergen | Non-Toxin | Positive |
| | **H2-IAb** | NLEDFQVAYTTPLLF | 0.9151 | Antigen | Non-allergen | Non-Toxin | Positive |
| | **H2-IAb** | LGEFSPGRISRTMMH | 0.7394 | Antigen | Non-allergen | Non-Toxin | Positive |
| | **H2-IAb** | PIGWNGYGAHPAALN | 0.8943 | Antigen | Non-allergen | Non-Toxin | Positive |
| | **VP5** | | | | | | |
| | **H2-IAb** | EEVPLIGAGMATAVA | 0.5161 | Antigen | Non-allergen | Non-Toxin | Positive |

**Supplementary Table 6.** List of finalized CD8+ and CD4+ T cell epitopes for the design of pan-BTV multi-epitope bovine vaccine.

| | Allele | Epitope | Antigenicity score | Antigenicty | Allergenicity | Toxicity | |
|---|---|---|---|---|---|---|---|
| | **VP1** | | | | | | |
| **Mouse CD8+ T cell epitopes** | BoLA-6:01301 | KMRMREEDI | 1.7662 | Antigen | Non-allergen | Non-Toxin | |
| | BoLA-6:01301 | IMISKLDEI | 0.5623 | Antigen | Non-allergen | Non-Toxin | |
| | BoLA-1:02301 | TKMSFFSKM | 1.1372 | Antigen | Non-allergen | Non-Toxin | |
| | BoLA-6:01301 | KINSRIKAL | 1.1637 | Antigen | Non-allergen | Non-Toxin | |
| | BoLA-6:01301 | ILMLKATFI | 1.1657 | Antigen | Non-allergen | Non-Toxin | |
| | BoLA-6:01301 | VMLRAALGL | 0.6255 | Antigen | Non-allergen | Non-Toxin | |
| | BoLA-1:02301 | KKYNSVELY | 0.5682 | Antigen | Non-allergen | Non-Toxin | |
| | BoLA-1:02301 | HKKYNSVEL | 1.1492 | Antigen | Non-allergen | Non-Toxin | |
| | BoLA-6:01301 | KLGSHDRRL | 1.5803 | Antigen | Non-allergen | Non-Toxin | |
| | **VP7** | | | | | | |
| | BoLA-6:01301 | TLQEARIVL | 0.5255 | Antigen | Non-allergen | Non-Toxin | |
| | BoLA-6:01301 | VQIQVVFYI | 1.0097 | Antigen | Non-allergen | Non-Toxin | |
| | **Allele** | **Epitope** | **Antigenicity score** | **Antigenicit y** | **Allergenicity** | **Toxicit y** | **IFNg inducing** |
| | **VP1** | | | | | | |
| | BoLA-DRB3_0101 | QMVMLRAALGLPK KK | 0.7323 | Antigen | Non-allergen | Non-Toxin | Positive |
| | BoLA-DRB3_1101 | ADPIVVLQSTAGDD R | 0.5582 | Antigen | Non-allergen | Non-Toxin | Positive |
| | **VP5** | | | | | | |

| | | | | | | | |
|---|---|---|---|---|---|---|---|
| **Mouse CD4+ T cell epitopes** | BoLA-DRB3_1201 | SIDLVHYEDLTAHAH | 1.0504 | Antigen | Non-allergen | Non-toxin | Positive |
| | **VP7** | | | | | | |
| | BoLA-DRB3_1001 | PMMIYLVWRRIENFA | 0.7744 | Antigen | Non-allergen | Non-Toxin | Positive |

**Supplementary Table 7.** Ramachandran plot analysis of the designed vaccine constructs.

| | **mVac-β-def** | **mVac-50s R** | **bVac-β-def** | **bVac-50s R** |
|---|---|---|---|---|
| **Most favoured** | 90.7 % | 93.3 % | 95.9 % | 97 % |
| **Additionally allowed** | 5.9 % | 5.5 % | 3.6 % | 2.6 % |
| **Generally allowed** | 1.5 % | 0.3 % | 0.0 | 0.0 % |
| **Disallowed** | 1.9 % | 0.9 % | 0.5 % | 0.4 % |
